# Leveraging publicly available datasets and machine learning approaches for predicting the health benefits of fermented foods

**DOI:** 10.64898/2026.03.05.709865

**Authors:** Elizabeth A. McDaniel, Chantle Edillor, Matthew Schertler, Rachel Dutton

## Abstract

Fermented foods are an ancient, near universal component of human dietary culture and are increasingly recognized for their health benefits. Bioactive peptides and biosynthetic gene clusters (BGCs) produced by microbes during fermentation have been shown to be key mediators of human health benefits, such as ACE inhibitors and antibacterial bacteriocins. To broadly map this potential, we leveraged the growing abundance of publicly available fermented food datasets alongside recent advances in machine learning models for bioactivity prediction. We collected, curated, and re-analyzed multiple publicly available multi-omics datasets from diverse fermented foods, enabling cross-food comparisons of microbial and molecular profiles. Our analyses include profiling a curated database of ∼1,300 species-representative genomes across hundreds of fermented food metagenomes, predicting genome-encoded BGCs and peptides from 11,500 bacterial genomes, and applying machine learning classification models to predict the bioactivity of thousands of genome-encoded and peptidomics-detected peptides. These models predict 17 different bioactivities, providing novel, testable predictions for downstream experimental characterization. Most importantly, all curated resources, underlying computational tools, and resulting datasets are publicly available to the community, with an emphasis on creating tools and resources that are user-friendly and empower the community to generate more efficient and predictive models for future fermented foods research. A descriptive list of all generated resources is available below, with all computational tools available on GitHub at https://github.com/MicrocosmFoods and all raw files and datasets available on Zenodo at https://zenodo.org/communities/microcosmfoods/.

**Resources:**

- [Dataset](https://zenodo.org/records/15588452) with ∼13,500 genomes from ∼3,000 different samples representing 150 foods and 50 countries
- [Species-representative dataset](https://seqhub.org/rachel-dutton/collections/microcosm-fermented-food-genomes) with ∼1,300 genomes based on 95% ANI, with full functional annotations available as an explorable collection on SeqHub
- [Strain-level representative dataset](https://kbase.us/n/218406/47/) with ∼4,300 genomes based on 99% ANI, available as a free and publicly available narrative on KBase for users to run their own analyses with
- Workflows for profiling metagenomic samples, performing functional annotation on bacterial genomes, and predicting peptide bioactivity using ML models
- [Predicted biosynthetic gene clusters and peptides](https://zenodo.org/records/15858778) for ∼11,500 bacterial genomes
- [Predicted bioactivities of peptides](https://zenodo.org/records/18750850) from ∼11,500 bacterial genomes and 5 proteomics datasets of fermented foods

## Background

Fermented foods are produced through the microbial transformation and enzymatic conversion of food components, where beneficial microbes convert sugars and other compounds into acids, alcohols, and other metabolites ^1,2^. These compounds act as preservatives such as organic acids that lower pH and bacteriocins that exhibit specific activity against foodborne microbial pathogens ^3,4^. Outside of food safety, fermentation can also enhance the nutrient availability of food through releasing or synthesizing essential vitamins such as vitamins and minerals ^5^. Additionally, fermentation contributes to sensory qualities and novel flavors of the food substrate. This ancient process spans nearly every culture globally and has resulted in diverse fermentation practices and foods, from dairy fermentations such as yogurt, cheese, and milk kefir to plant-based fermentations such as sauerkraut, kimchi, and sourdough bread.

Bioactive molecules produced by microbes during the fermentation process such as organic acids, peptides, and small molecules have been increasingly recognized as potential mediators of human immune and metabolic health ^1^. Similar to the metabolism of microbes in the gut, food fermentation produces human-relevant compounds such as short-chain fatty acids like acetate, butyrate, and propionate that interact (either directly or indirectly via the gut microbiome) with specific tissues and strengthen gut barrier integrity, modulate metabolic function, and regulate immune responses ^6,7^. D-phenyllactic acid (D-PLA) is produced by lactic acid bacteria and found in foods such as sauerkraut and kimchi, and activates human hydroxycarboxylic acid receptor 3 (HCA3), which is hypothesized to play a role in immunomodulation ^8–10^. The consumption of various fermented foods such as yogurt, kimchi, and kombucha has been associated with decreased markers of inflammation and increased gut microbiome diversity ^11^.

Recent large-scale metagenomic surveys have generated a wealth of microbial genome data from diverse fermented foods. This abundance of data presents an opportunity to move beyond single-food or single-molecule studies and systematically map the potential for bioactive molecules across the entire fermented food landscape. However, fully leveraging this data requires costly sophisticated computational approaches and accessible resources. Machine learning classification models, which have seen significant development in the bioactive peptide and BGC spaces, offer a powerful way to increase the prediction and hypothesis space, enabling researchers to move from descriptive characterizations to testable predictions of what molecules interact with human biology. We decided to focus our efforts on peptide bioactivity prediction, since there is a better availability of user-friendly tools and established ML models for many bioactivities, and the relative ease of downstream experimental validation due to cheap peptide synthesis technologies. A graphical overview of the study and the underlying computational workflow architecture is summarized in Figure 1.

**Figure 1.**
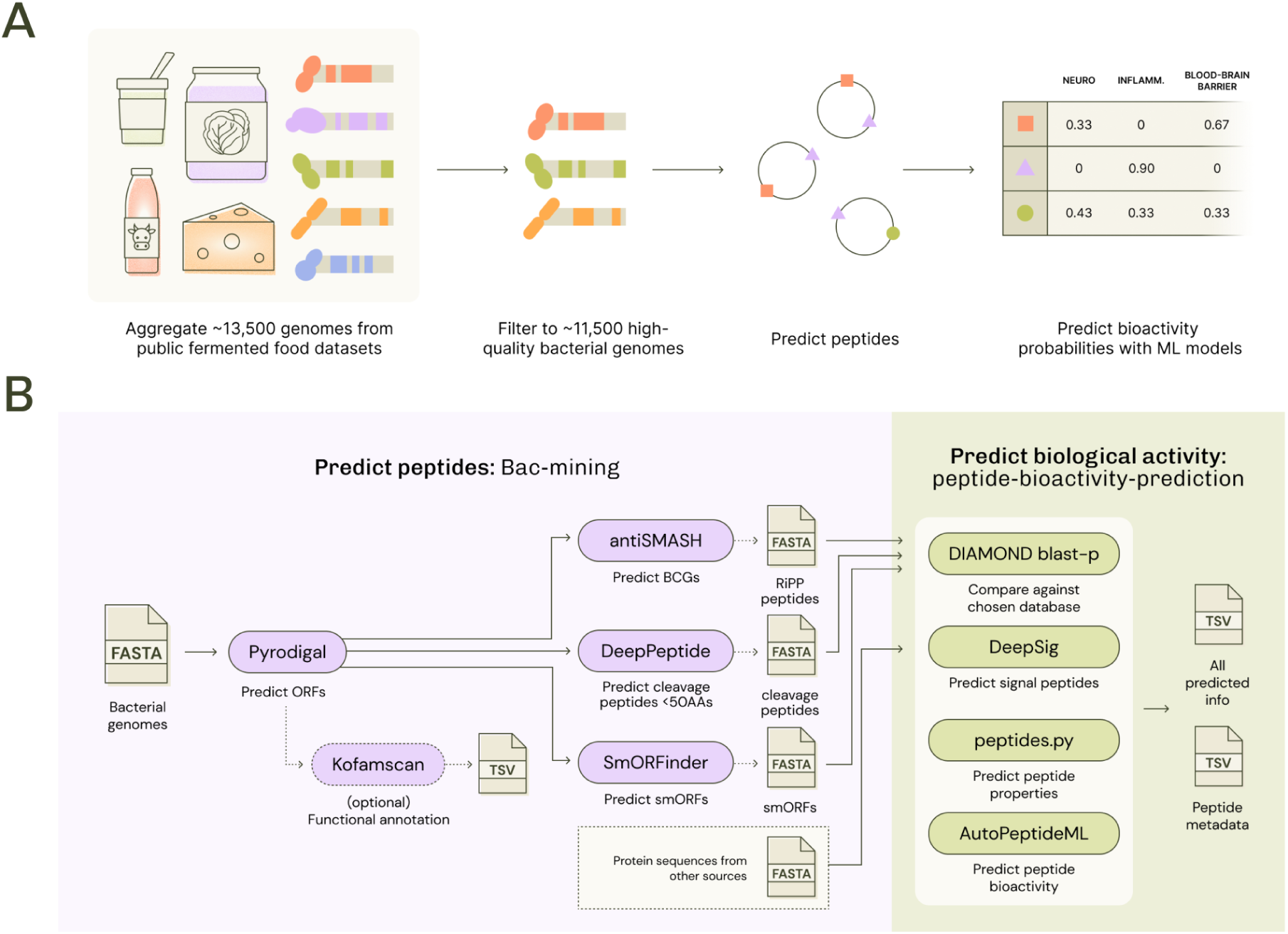
Peptide Mining Overview and Computational Workflow Architecture. A) Strategy for mining peptides from fermented food associated bacterial genomes. The entire fermented food microbial genome dataset containing ∼13,500 genomes was filtered to only include ∼11,500 high-quality bacterial genomes, where quality is defined in the methods. Peptides were predicted from open reading frames and used for predicting bioactivity with 17 machine learning models. B) Computational workflow architecture for bac-mining workflow for bacterial genome annotation and peptide-bioactivity-prediction workflow for predicting peptide characteristics and bioactivity. For the bac-mining workflow, the input is a set of bacterial genomes in FASTA format, and the output are annotations from antiSMASH for predicting biosynthetic gene clusters, DeepPeptide for predicting cleavage peptides smaller than 50 AAs, and smORFinder for predicting small ORF peptides. Optionally Kofamscan for full bacterial genome annotation can be run as well. The input for the peptide-bioactivity-prediction workflow is a set of peptide sequences in FASTA format, which can come from the output of predicting genome-encoded peptides or directly from a proteomics experiment. The output is a TSV file containing for each input peptide sequence the predicted physicochemical characteristics, signal peptide predictions, bioactivity predictions, and DIAMOND blast-p hits to an input database of the users choice.

In this work, we have collected, curated, and re-analyzed multiple types of datasets from fermented foods. This encompasses curated microbial genomic datasets, predicted functional annotations of these genomes, and bioactive peptide predictions. All results and resources are associated with publicly available datasets, workflows, or links to other usable platforms for the community to explore, reuse, and build upon.

## Results and Discussion

### Curated set of ∼13,500 genomes from diverse fermented foods with associated metadata

We first sought to collect and curate microbial genomes from diverse fermented foods with associated metadata. Numerous studies have recently released publicly curated isolate genomes and metagenome-assembled genomes (MAGs) from fermented foods, including two large meta-studies of microbial genomes from food samples. MiFoDB (Microbial Food DataBase) is a collection of 675 genomes of bacteria, yeast, fungi, and common fermented food substrates curated from metagenomic sequencing of 90 fermented foods and other publicly available genomes ^12^. This database was used in coordination with a workflow for tracking strain-level dynamics in different fermented foods. The curatedFoodMetagenomicData (cFMD) database is a collection of nearly 11,000 microbial genomes from 2500 food metagenomes and was used for profiling categories of microbial genomes in foods and probing the overlap of microbial species in human versus food metagenomes ^13^. A subset of this database has already been reused to probe the functional profiles of acetic-acid bacteria from fermented food samples ^14^. We hypothesized that creating a well-curated microbial genomes database primarily sourced from these two databases and a handful of other studies would enable broad community use for downstream specific research questions. Additionally, we carefully curated the associated metadata for fermented food-relevant information to easily allow researchers to link their predictions to metadata features.

The curated set of ∼13,500 genomes, including reference isolates and metagenome-assembled genomes, spans ∼3,000 different samples, over 150 different foods, and originating from at least 50 different countries (Figure 2). We decided to dereplicate these genomes at both 95% average nucleotide identity (ANI) and 99% ANI to represent species-level and “strain-level” representatives, since dereplicating at species-level would help remove redundancy, but dereplicating at strain-level would preserve potentially important strain-level bioactivities or differences between genomes across food substrates.

**Figure 2.**
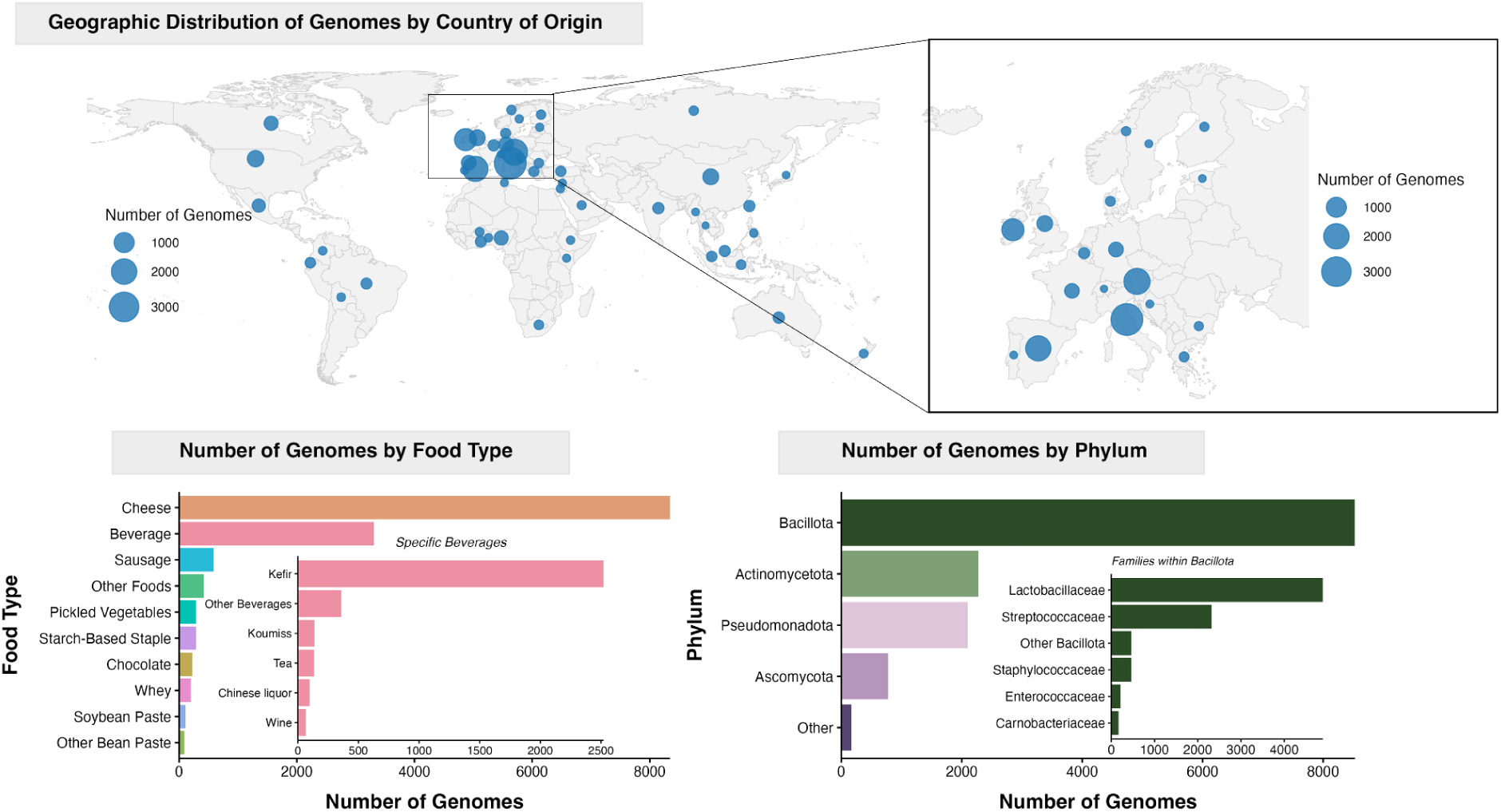
Overview of the Fermented Food Microbial Genomes Database. For each genome, we curated metadata including i) the country of origin of the metagenomic sample from which the genome was assembled (when reliably available), ii) the sample’s food type, and iii) the genome’s phylum assigned by GTDB-tk v2.4.1 with reference data r226. Subset panels include a Europe zoom-in to distinguish genomes originating from European countries; a breakdown of “beverage”-classified genomes, and genome counts by family within Bacillota.

We have uploaded the curated metadata and raw genomes for the ∼4,000 strain-level representatives as a narrative to KBase^15–17^, where users can freely explore this dataset further by subsetting into genomes of interest into their own narratives, run analyses such as metabolic predictions and pangenomic comparisons, and incorporate their own genomes in addition to the fermented foods microbial genomes collection.

### Profiling species-representative genomes against select metagenomic samples

To test if our species-representative database of microbial genomes captures the diversity of fermented foods samples, we profiled our database against a set of publicly available fermented food metagenomic samples from Carlino et al.^13^ (Figure 3). We selected a maximum of 10 samples from each fermented food type so as not to overrepresent highly common food types such as dairy samples, resulting in 366 samples for profiling. All input samples had 99-100% mapping rates against our species-representative database, giving us confidence that we have a well covered database for the fermented food metagenomic samples we profiled.

**Figure 3.**
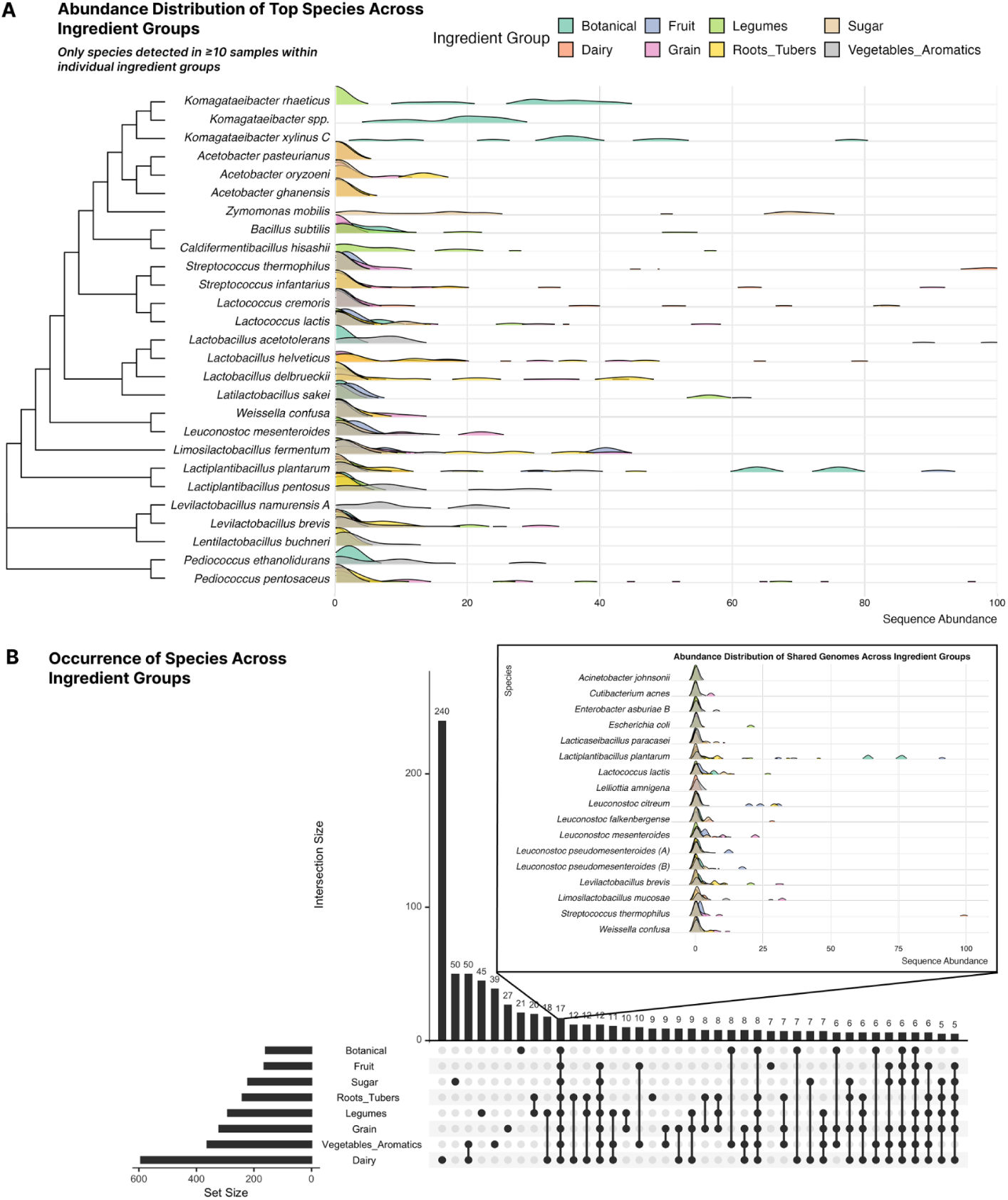
Metagenomic profiling of the species-representative genomes among select fermented food samples. A) Abundance distribution of bacterial species that are present in at least 10 samples within an individual ingredient group. The y-axis is the frequency of samples within that ingredient group that exhibit the abundance pattern (x-axis) of that particular species. Species are shown based on a concatenated phylogenetic tree of core ribosomal markers. Botanical based foods include tea and mustard plants. Roots and tubers foods include turnip and cassava. Sugar based foods include water kefir and pulque, an agave based drink. Aromatics include ginger. B) Upset plot of the occurrence of species across select ingredient groups. The x-axis or set size shows the total number of genomes detected in that ingredient group. The y-axis set size is the number of genomes found exclusively in the individual or intersections of ingredient groups.

To understand how similar or diverging the microbial composition of different fermented foods may be, we analyzed the abundance distribution of top species detected in the public metagenomes dataset. We first analyzed the abundance distribution of genomes detected in at least 10 samples within individual ingredient groups (Figure 3A). Most genomes that are detected in multiple samples and across ingredient groups are present at lower abundances, usually 1-5% of the total mapped metagenomic reads. This is also represented in an analysis of the overlap of detected genomes across ingredient groups (Figure 3B), where genomes detected in at least 10 samples across all ingredient groups are mostly present at low abundance in all samples. Genomes detected at higher abundances such as greater than 50% of mapped reads are exclusive to specific ingredient groups, such as *Zymomonas mobilis* in sugar-based foods, *Lactiplantibacillus plantarum* in botanical-based foods, and *Streptococcus thermophilus* in dairy-based foods.

### Predicted biosynthetic gene clusters and peptides from a subset of bacterial genomes

We were interested in predicting biosynthetic gene clusters and peptides from our curated genomes dataset since these types of molecules from bacteria often exhibit some type of activity relevant for human health, ranging from antimicrobial to immunomodulatory activity. For example, biosynthetic gene clusters predicted from fermented food microbial genomes were previously used to predict antibiotic and antifungal activity using machine learning classification methods^19^. Large-scale peptide prediction efforts from bacterial genomes paired with machine learning approaches and subsequent experimental validation have also yielded multiple novel antibiotic and immunomodulatory peptide candidates ^20,21^. Therefore, we hypothesized this would be a valuable dataset for the community ranging from researchers interested in the basic biology of biosynthetic gene clusters to those experimentally validating machine learning predictions of peptide bioactivity.

We selected a subset of the ∼13,500 genomes from our curated dataset, selecting only bacterial genomes with greater than 50% completion, less than 10% redundancy, and an N50 greater than 8000. This resulted in a total of ∼11,500 bacterial genomes to predict biosynthetic gene clusters and peptides from. We predicted all biosynthetic gene cluster types using antiSMASH, cleavage peptides with DeepPeptide, and small ORF peptides (smORFs) with smORFinder. We did not dereplicate any of the genomes or de-duplicate any of the predicted molecules by sequence similarity, and show total counts across all molecule types. We made this decision because we hypothesized that the presence of BGCs and peptides could be highly strain-dependent, and therefore to maximize exploration potential of this dataset we made these predictions for all genomes and left downstream filtering and dereplication for future users.

We visualized the counts of predicted molecule types across the top five fermented foods - cheese, kefir, salami, chocolate and sourdough (Figure 4). We also analyzed for the three top phyla for genomes within these fermented food categories the counts of molecule types. This included *Actinomycetota, Bacillota,* and *Pseudomonadota*. Within these food categories, there are more than three times as many genomes belonging to *Bacillota* than *Actinomycetota*. Relative to the total number of genomes in these phyla and food categories, many of the biosynthetic gene clusters are predicted from *Actinomycetota.* However, a larger count of RiPP-like molecules are predicted from *Bacillota.* Additionally, the counts of cleavage peptides and small ORFs (smORFs) are also relatively higher in *Bacillota* than *Actinomycetota*.

**Figure 4.**
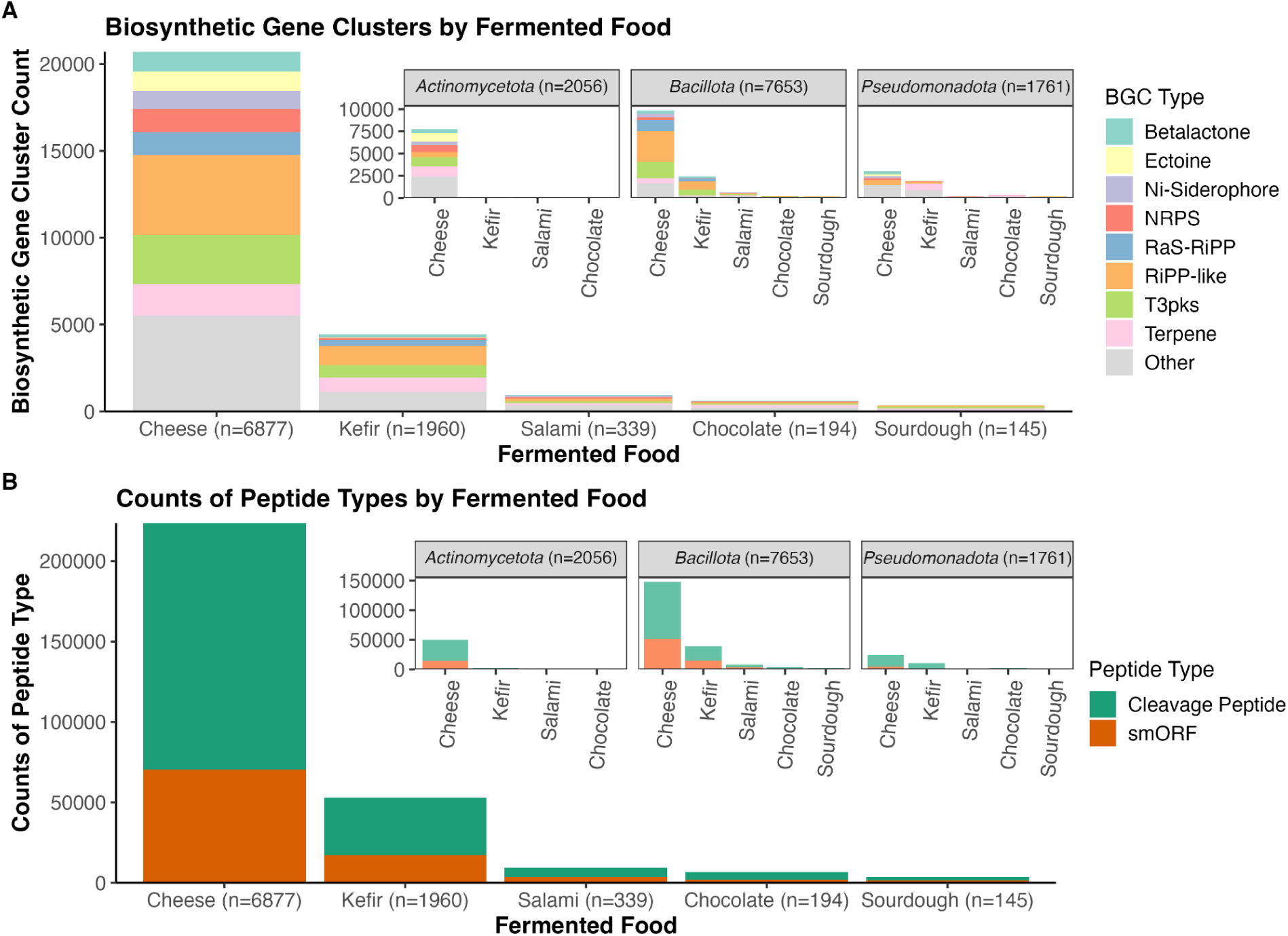
Counts of Biosynthetic Gene Clusters and Peptides across Top Fermented Foods. Counts of major types of A) biosynthetic gene clusters predicted with antiSMASH and B) counts of cleavage peptides predicted by DeepPeptide and small ORFs (smORFs) predicted by smORFinder. Counts are summarized for the top five fermented food categories in the dataset, where the number of bacterial genomes represented in that food category is listed on the x-axis. Inset plots show counts of each molecule type for the three largest phyla represented in the dataset across the five fermented food types, where the number of genomes in that phylum are represented in the facet labels.

### Predicted bioactivity of peptides from bacterial genomes and fermented food proteomics experiments

Fermented foods have long been known to be a source of bioactive peptides, with the most well-known being bacteriocins produced by lactic acid bacteria that kill contaminating organisms and contribute to food safety ^22^. However, many peptides from fermented foods have been experimentally shown to have other bioactive properties outside of antibiotic activity, such as ACE-inhibitor effects, antioxidative properties, and immunomodulatory activities ^23–25^. Most commonly, bioactive peptides have been identified from fermented foods through proteomics experiments. These sequences are compared to databases of peptides with known bioactivity and only those with 100% sequence identity are assigned that bioactivity^24^. However, applying machine learning classification models to large pools of genome-encoded peptides or peptides detected from high-throughput proteomics experiments can substantially increase the pool of candidate peptides with diverse bioactivities to select from for downstream experimental validation^21,26^. Additionally, using the wealth of genomic data curated from metagenomic surveys of fermented foods, bioactive peptides can also be predicted *in silico* from genome-encoded peptide predictions.

To identify putatively bioactive peptides from fermented foods, we applied machine learning classification models of 17 bioactivities to genome-encoded peptides predicted from our bacterial genome collection described above and 5 proteomics datasets of fermented foods. The 5 proteomics datasets include samples from sheep milk kefir ^27^, four different cheeses ^24^, fermented cocoa beans ^28^, fermented donkey milk ^29^, and kefir grains ^30^. We inputted the set of all predicted peptides from the bacterial genomes and the raw peptide sequences from the proteomics experiments through a computational workflow that predicts physiochemical properties of the peptides, compares the sequences to a subset of sequences in the Peptipedia database that have experimentally verified activity for select bioactivities, and predicts bioactivities with the 17 machine learning models. Metrics for each bioactivity model are provided in Table 1 in the methods.

**Table 1.** Peptide bioactivity machine learning model performance metrics. Performance metrics for the custom anti-inflammatory model reported in the output of AutoPeptideML, as well as metrics for the 16 models generated through AutoPeptideML. ANIF = Antiinflammatory, AB = Antibiotic, ACE = ACE-inhibitor, ACP = Anti-cancer peptide, AF = Antifungal, AMAP = Antimalarial peptide, AMP = Antimicrobial, AOX = Antioxidant, APP = Antiparasitic, AV = Antiviral, BBP = Blood-brain penetrating, DPPIV = DPP IV inhibitor, MRSA = Anti-MRSA, NEURO = Neuropeptide, QS = Quorum sensing, TOX = toxic, TTCA = Tumor T-cell antigens.

| Model | Specificity | Sensitivity | Accuracy | MCC | AUC/ROC | Source |
| --- | --- | --- | --- | --- | --- | --- |
| ANIF | 0.75 | 0.84 | 0.73 | 0.40 | 0.76 | This study |
| AB_1 | 0.81 | 0.61 | 0.73 | 0.47 | 0.79 | AutoPeptideML |
| ACE_1 | 0.76 | 0.79 | 0.77 | 0.54 | 0.83 | AutoPeptideML |
| ACP_1 | 0.77 | 0.42 | 0.65 | 0.33 | 0.69 | AutoPeptideML |
| AF_1 | 0.68 | 0.42 | 0.61 | 0.24 | 0.66 | AutoPeptideML |
| AMAP_1 | 0.62 | 0.78 | 0.65 | 0.32 | 0.75 | AutoPeptideML |
| AMP_1 | 0.66 | 0.60 | 0.64 | 0.29 | 0.73 | AutoPeptideML |
| AOX_1 | 0.57 | 0.58 | 0.57 | 0.15 | 0.65 | AutoPeptideML |
| APP_1 | 0.63 | 0.66 | 0.63 | 0.27 | 0.69 | AutoPeptideML |
| AV_1 | 0.76 | 0.78 | 0.77 | 0.64 | 0.84 | AutoPeptideML |
| BBP_1 | 0.52 | 0.54 | 0.51 | 0.02 | 0.53 | AutoPeptideML |
| DPPIV_1 | 0.75 | 0.79 | 0.76 | 0.53 | 0.83 | AutoPeptideML |
| MRSA_1 | 0.67 | 0.55 | 0.64 | 0.29 | 0.78 | AutoPeptideML |
| NEURO_1 | 0.76 | 0.89 | 0.80 | 0.62 | 0.90 | AutoPeptideML |
| QS_1 | 0.70 | 0.86 | 0.75 | 0.51 | 0.89 | AutoPeptideML |
| TOX_1 | 0.61 | 0.76 | 0.63 | 0.27 | 0.74 | AutoPeptideML |
| TTCA_1 | 0.83 | 0.92 | 0.86 | 0.73 | 0.93 | AutoPeptideML |

To analyze the peptide predictions, we clustered the genome-encoded and proteomics peptide samples separately at 100% sequence identity to reduce redundancy of identical peptide sequences from the analysis. We then filtered the results to remove peptides that had a toxic bioactivity prediction with probability > 0.5 and kept a bioactivity designation for a peptide if the probability for the bioactivity was > 0.9. For both peptide datasets we summarized the counts of predicted bioactivity categories, the most frequent intersections of bioactivity categories, and cases where we detected peptide sequences in our datasets that were identical to experimentally verified peptides in the literature as curated in Peptipedia.

For the genome-encoded peptides dataset, we started with a total of 674,113 peptide sequences reduced to 228,956 non-redundant sequences and then analyzed the bioactivities of 63,995 sequences after filtering out non-toxic peptides and keeping bioactivities with probability > 0.9 (Figure 5). The top bioactivity categories of this pool of peptides were neuropeptide, blood-brain barrier penetrating, antioxidative, and antiviral (Figure 5A), and the largest intersections of bioactivity categories being different combinations of these four activities (Figure 5B).

**Figure 5.**
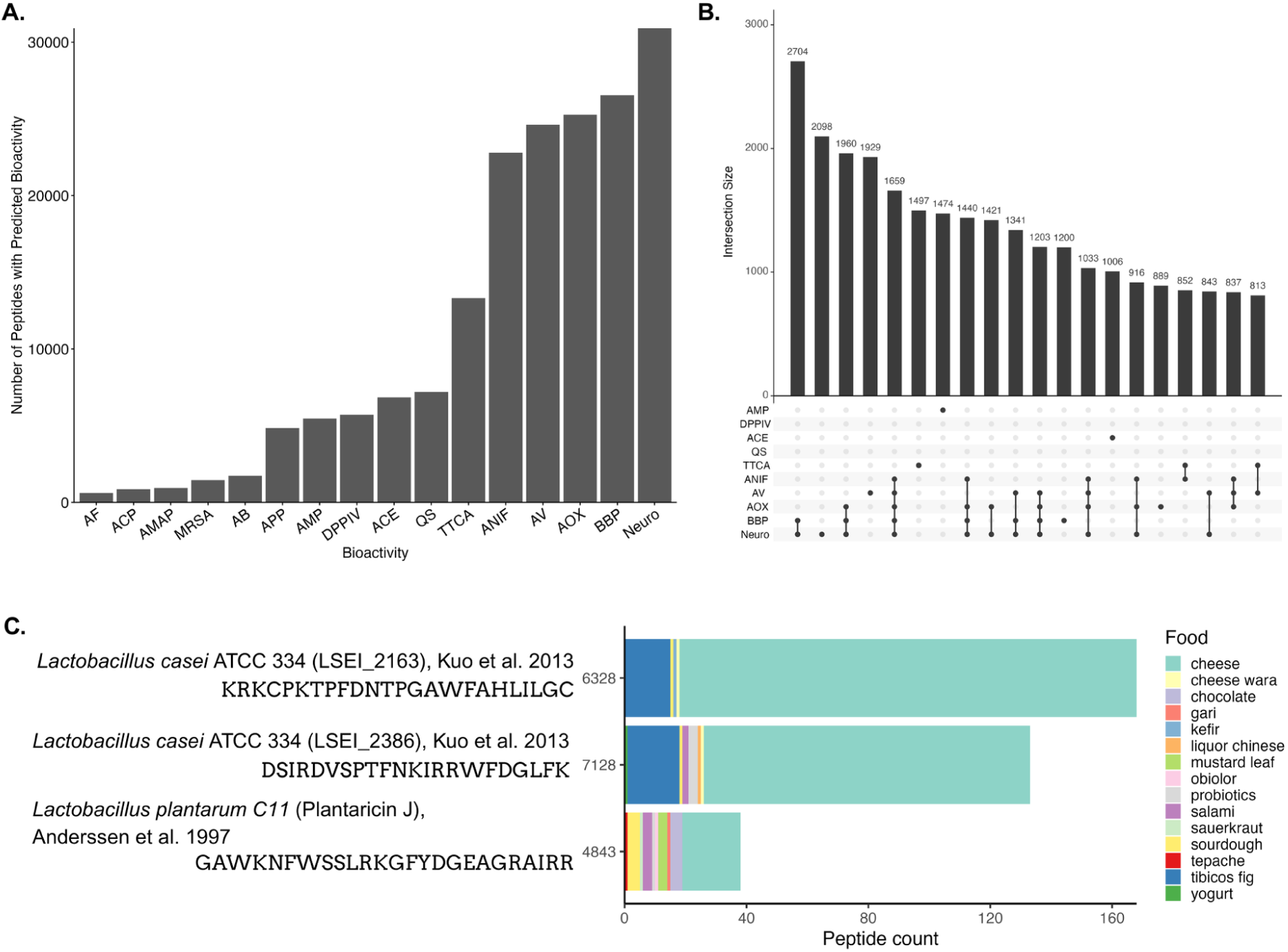
Overview of predicted genome-encoded bioactive peptides. For all results, peptides were clustered at 100% sequence identity to reduce redundancy and only cluster-representatives are plotted. To count a peptide as having that predicted bioactivity, we removed peptides with a toxic bioactivity probability > 0.5, and kept a predicted bioactivity if the probability was above 0.90. A) Raw counts of peptides and predicted bioactivities. B) Intersections of counts of peptides with predicted bioactivities. C) Select DIAMOND blastp hits of predicted genome encoded peptides to sequences in the Peptipedia database with experimentally verified antibiotic activity. Each of these three examples are experimentally verified bacteriocin peptides with antibiotic activity. The peptide count represents the number of peptides in the cluster and their food origin that the genome was assembled from. The x-axis code represents the Peptipedia database ID, with additional information about the peptide sequence, species it was originally studied in, and the study. Bioactivity abbreviations: ANIF = Antiinflammatory, AB = Antibiotic, ACE = ACE-inhibitor, ACP = Anti-cancer peptide, AF = Antifungal, AMAP = Antimalarial peptide, AMP = Antimicrobial, AOX = Antioxidant, APP = Antiparasitic, AV = Antiviral, BBP = Blood-brain penetrating, DPPIV = DPP IV inhibitor, MRSA = Anti-MRSA, NEURO = Neuropeptide, QS = Quorum sensing, TOX = toxic, TTCA = Tumor T-cell antigens.

To test if we could detect peptides in this dataset that already had evidence of an experimentally validated bioactivity, we compared our peptide sequences to those in the Peptipedia database labelled with some amount of experimental evidence either in a database or documented in the literature. For the genome-encoded peptides, the largest amount of hits in our dataset against experimentally verified peptides in Peptipedia with 100% sequence identity and length were to those with antibiotic activity, specifically bacteriocins identified in *Lactobacillus* isolates ^22,31^ (Figure 5C). Two peptide sequences identified in our dataset with experimentally verified bacteriocin activity were most frequently found in cheese samples, but also tibicos fig/water kefir (Figure 5C).

To test whether our machine learning models could accurately predict positive examples of peptides with antimicrobial activity in our genome-encoded peptides dataset, we compared sequences with a hit to a Peptipedia sequence with experimentally verified antimicrobial activity to the predicted probability score for the antibiotic and antimicrobial models (Supplementary Table 1). For these sequences, there are oftentimes instances where the antibiotic and antimicrobial models don’t closely agree, but these models were trained on different benchmark datasets. Overall, there are only a few instances for these experimentally verified sequences where neither model has a predicted probability above 0.5, representing sequences that our models miss.

For the proteomics peptides dataset, we started with a total of 114,325 peptide sequences reduced to 11,869 non-redundant sequences and analyzed the bioactivities of 4,012 peptides after filtering for non-toxic peptides and keeping bioactivities with probability > 0.9 (Figure 6). The top predicted bioactivity categories for this pool of peptides were DPP-IV inhibitors, antioxidative activity, tumor T-cell antigens, and ACE-inhibitors (Figure 6A). The top intersections of bioactivity categories were DPP-IV inhibitors, antioxidative, and ACE-inhibitors, and DPP-IV and ACE-inhibitors (Figure 6B).

**Figure 6.**
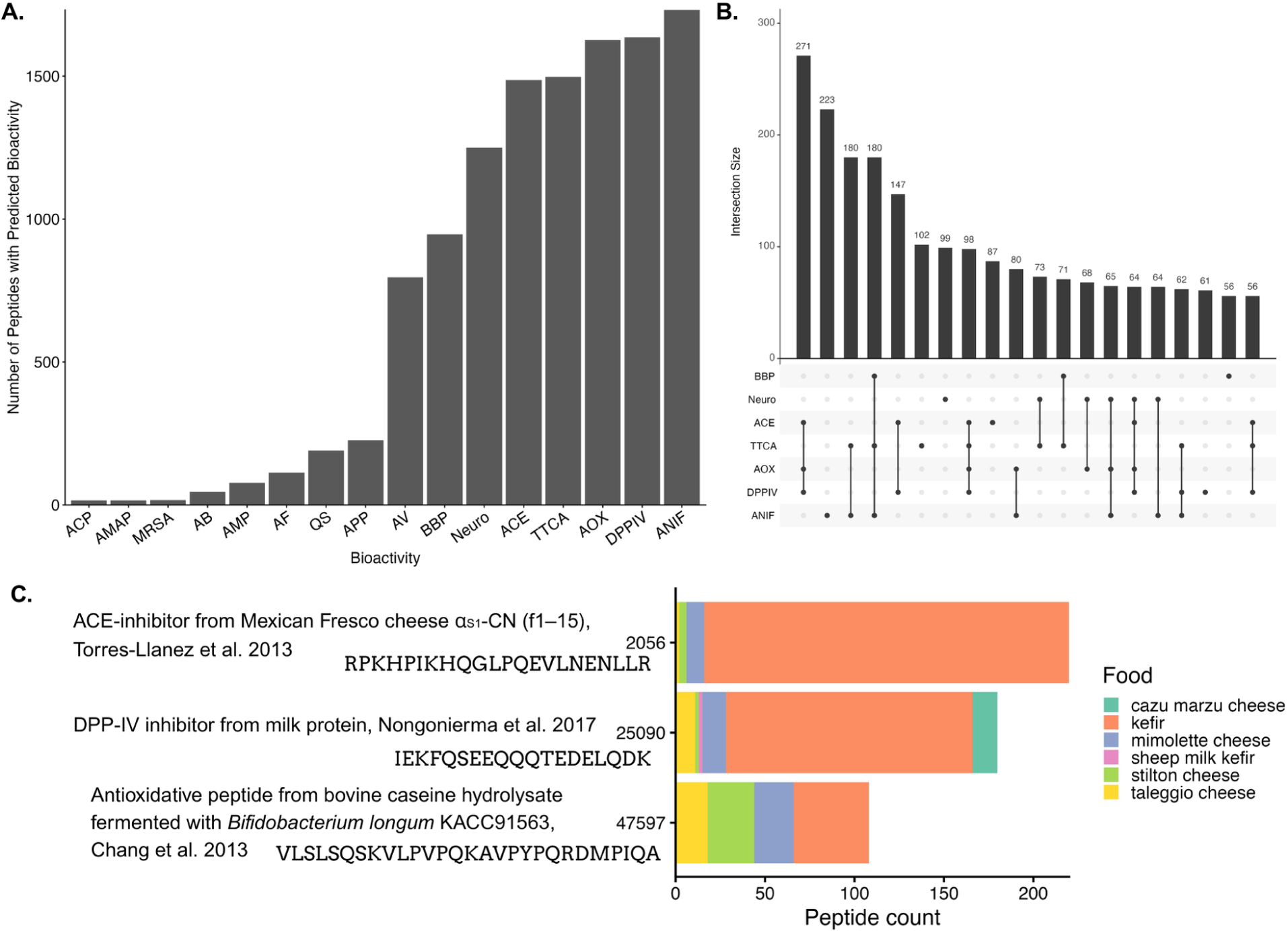
Overview of predicted bioactive peptides from fermented foods proteomics experiments. For all results, peptides were clustered at 100% sequence identity to reduce redundancy and only cluster-representatives are plotted. A) Raw counts of peptides and predicted bioactivities. B) Intersections of counts of peptides with predicted bioactivities. C) Select DIAMOND blastp hits of peptides to sequences in the Peptipedia database with experimentally verified activities. The peptide count represents the number of peptides in the cluster and the food proteomics sample the peptide was originally from. The x-axis represents the Peptipedia database ID, with additional information about the peptide sequence, type of sample the peptide was originally found and tested, and the specific bioactivity that study confirmed activity for. Bioactivity abbreviations: ANIF = Antiinflammatory, AB = Antibiotic, ACE = ACE-inhibitor, ACP = Anti-cancer peptide, AF = Antifungal, AMAP = Antimalarial peptide, AMP = Antimicrobial, AOX = Antioxidant, APP = Antiparasitic, AV = Antiviral, BBP = Blood-brain penetrating, DPPIV = DPP IV inhibitor, MRSA = Anti-MRSA, NEURO = Neuropeptide, QS = Quorum sensing, TOX = toxic, TTCA = Tumor T-cell antigens.

To test if we could detect peptides from the proteomics experiments that already had evidence of an experimentally validated bioactivity, we compared our peptide sequences to those in the Peptipedia database labelled with some amount of experimental evidence either in a database or documented in the literature. The peptides with the greatest amount of hits to a peptide sequence in the Peptipedia database with 100% sequence identity and length had experimentally verified bioactivities for ACE-inhibitor activity, DPP-IV inhibitor activity, and antioxidative properties ^23,32,33^ (Figure 6C). The hits to the peptides with these activities were most frequently identified in the kefir samples, as this sample had the highest amount of starting peptide sequences compared to the other samples, and different cheese samples. Since the proteomics samples are collected from the fermented food itself, peptide sequences could come from hydrolyzed substrate protein produced during fermentation, and not derived directly from peptides encoded in the microbial genomes, such as identifying hits to the sequences with ACE-inhibitor and DPP-IV inhibitor activity from Mexican Fresco cheese and milk protein, respectively.

To test whether our machine learning models could accurately predict positive examples of peptides with anti-inflammatory or ACE-inhibitor activity in the proteomics samples, we compared sequences with a hit to a Peptipedia sequence with these experimentally verified activities to the predicted probability score for the corresponding machine learning models (Supplementary Tables 2 and 3). We only had 4 hits to experimentally validated sequences with anti-inflammatory activity (Supplementary Table 2), and all but one of these sequences had a greater than 0.5 probability score predicted with our custom anti-inflammatory classification model. One of the sequences is present nearly 40 times across the proteomics dataset. When we performed blast-p searches for this peptide, we got hits to part of the alpha-casein protein. For hits to sequences with experimentally validated ACE-inhibitor activity the results were more varied with predicted probabilities by the model, with 6 sequences predicted with a less than 0.5 probability score. These peptide hits were also to parts of the alpha-casein protein, which has confirmed ACE-inhibitory activity. Overall, we were able to validate our machine learning approach with validated experimental examples from the literature, but there is an enormous need to further experimentally validate the bioactivity predictions of these peptides, especially for activities that have not normally been tested in fermented foods.

## Conclusions

In this work, we present a comprehensive framework for re-using publicly available data and applying machine learning models to generate predictions for downstream experimental characterization. Our central goal was to systematically map the broad diversity and potential for bioactive peptides across the fermented food landscape by leveraging the abundance of publicly available data on fermented foods, namely genomes generated from large-scale metagenomic surveys. By applying ML models to predict the bioactivity of thousands of peptides, our approach significantly expands the prediction and hypothesis space, moving beyond descriptive characterization of species presence to propose novel, testable mechanisms for how fermented food-derived molecules may interact with human biology.

Through these analyses, we generated several key insights:

- We mapped the distribution of microbial species across various food types and identified both broad-range and food-specific keystone species.
- Applying 17 bioactivity machine learning classification models generated high-confidence predictions for over 600,000 genome-encoded peptides (top activities being neuropeptide, blood-brain barrier penetrating, antioxidative, and antiviral), and over 114,000 proteomics sourced peptides (top activities: antiinflammatory, DPP-IV inhibitor, antioxidative, tumor T-cell antigens, ACE-inhibitors)
- We validated our peptide bioactivity machine learning approach by identifying peptide sequences in both datasets that were identical to those with experimentally confirmed bioactivities in the literature, such as bacteriocins and ACE-inhibitors.

Importantly, this work provides testable predictions that need to be experimentally validated. There are several limitations to our *in silico* predictions of peptide bioactivity. First, most of our results are based on predicting peptides from genomic open reading frames, not experimentally validated peptide sequences like those obtained from proteomics approaches. For example, proteolytic peptide prediction tools may overestimate the actual peptide pool as they primarily simulate digestion with eukaryotic proteases, and therefore these might not be biologically real peptides. Second, many of the predicted peptides might not be stable in the human digestive system or be bioavailable. Follow-up experiments simulating digestive enzymatic processes will be needed to understand the relevance of these peptides in the context of human biology. And third, the accuracy of the bioactivity models are inherently limited by their training data. Models for predicting antibacterial and antifungal activity are likely more accurate than those for anti-inflammatory activity due to their larger training datasets stemming from ease of experimental characterization of these activities. Particularly for our custom anti-inflammatory activity model, we struggled to find high-quality, large training datasets. Most benchmark training datasets for anti-inflammatory peptide activity are extremely small and not as diverse compared to antibacterial peptide activity. We primarily focused on using public databases and tools for constructing our models such as creating training datasets from publicly available peptide sequences in Peptipedia and generating models with AutoPeptideML to demonstrate for the community what can be done with these existing tools out of the box. These tools, models, and predicted bioactivity data need to be evaluated with experimental validation and used for further training more confident models.

Importantly, given the caveats summarized above for our peptide bioactivity predictions, we’ve made all of our raw data, predictions, and models available for the community to build upon at different stages. All fermented food microbial genomes are available for researchers, whether to apply different peptide prediction tools or pursue other research interests. All of our raw peptide sequence predictions are available as well as the predicted physicochemical properties for the community to test existing models or build entirely new bioactivity models with. And finally, the raw model output for our custom anti-inflammatory model from AutoPeptideML is available for evaluation and comparison to other models.

More importantly, a core contribution of this work outside of the focus on bioactive peptides from fermented foods is the development of comprehensive, immediately usable, and publicly available resources and tools for the research community to build upon. This includes:

- [Dataset](https://zenodo.org/records/15588452) with ∼13,500 genomes from ∼3,000 different samples representing 150 foods and 50 countries
- [Strain-level representative dataset](https://kbase.us/n/218406/47/) with ∼4,300 genomes based on 99% ANI, available as a free and publicly available narrative on KBase for users to run their own analyses with
- [Species-representative dataset](https://seqhub.org/rachel-dutton/collections/microcosm-fermented-food-genomes) with ∼1,300 genomes based on 95% ANI, with full functional annotations available as an explorable collection on SeqHub
- Workflows for profiling metagenomic samples, performing functional annotation on bacterial genomes, and predicting peptide bioactivity using ML models
- [Predicted biosynthetic gene clusters and peptides](https://zenodo.org/records/15858778) for ∼11,500 bacterial genomes
- [Predicted bioactivities of peptides](https://zenodo.org/records/18750850) from ∼11,500 bacterial genomes and 5 proteomics datasets of fermented foods

In summary, this work starts to move from descriptive characterizations of fermented foods towards establishing a predictive framework for the field. By providing all curated data and tools openly, we aim to accelerate the discovery and validation of fermented food-derived molecules such as bioactive peptides for a more predictive and mechanistic understanding of how these foods interact with human biology.

## Methods

### Genome data collection and curation

We collected bacterial and eukaryotic genomes assembled from diverse fermented foods primarily from two sources, Carlino et al. 2024^13^ and Caffrey et al. 2025^12^, with a handful of MAGs also from Saak et al. 2023^34^, Rappaport et al. 2024^35^, and Du et al. 2023^19^. We curated metadata from all of these studies and genomes with a custom fermented food ontology that we based on the original sample description in the SRA run deposit and/or the original paper that the genome was produced through. We listed country of origin for each sample that we could find this information for, and for genomes that were coassembled from multiple samples we listed the range of SRA/ENA accessions that the genome could have originated from. We attempted to remove duplicate genomes and/or samples based on the SRA runs or study accessions. Genome quality information such as number of contigs, n50, etc. was added for each genome using QUAST v5.2.0^36^. Genomes were then dereplicated using dRep v3.6.2 ^37^ at 95% ANI and 99% ANI, resulting in ∼1300 species-level and ∼4300 strain-level representatives, respectively. The ∼1300 species-level genomes were assigned taxonomy using GTDB-tk v2.4.1 with reference data r226^38,39^. Code documenting how we curated the sets of genomes and associated metadata is available on GitHub at https://github.com/MicrocosmFoods/fermentedfood_mags_curation and https://github.com/MicrocosmFoods/fermentedfood_metadata_curation.

### Metagenomic sample profiling

To quantify the coverage of our genome database, we profiled a select set of metagenomic samples against our set of ∼1300 species-level representative genomes. We created a metagenome-containment-profiler workflow that uses sylph v0.8.1^40^ to quantify the containment of genomes in each metagenomic sample. By default sylph counts a genome as “present” if the ANI of the hit is 95%. Importantly the input reference database must be dereplicated at the species-representative or 95% ANI cutoff, as genomes that are more similar above this ANI threshold won’t be discriminated against or counted correctly using sylph. For all metagenome profiling runs we use our ∼1300 species-representative database. This resource is available as a Nextflow workflow and handles automating metagenomic sample and reference genome downloads, running sylph, and collating results, and is available on GitHub at https://github.com/MicrocosmFoods/metagenome-containment-profiler. Scripts for analyzing and visualizing the output files of this workflow are also available in the GitHub repository. A phylogenetic tree of the top most present bacterial genomes was created using metabolisHMM v 2.21^41^, using a core set of ribosomal protein markers ^42^. The tree was visualized using ggtree to place alongside the relative abundance plot of these top most present bacterial genomes.

### Genome mining and annotation workflow

We selected a subset of the ∼13,000 MAGs from the dataset, selecting only bacterial MAGs with >50% completion, <10% redundancy, and N50 > 8000. This resulted in a total of ∼11,500 bacterial genomes for mining molecules from. We created a Nextflow workflow bac-mining to perform several functional annotation steps. This includes predicting biosynthetic gene clusters, two peptide types - small ORFs (smORFs) and cleavage peptides, and optionally full functional annotation with Kofamscan using KEGG HMMs. We only performed full functional annotation with Kofamscan on the set of ∼1,100 species-representative bacterial genomes that passed the quality threshold, as it was cost prohibitive to run full functional annotation on all ∼11,500 genomes. The workflow is available on GitHub at https://github.com/MicrocosmFoods/bac-mining.

First, open reading frames are predicted from input genomes using Pyrodigal v3.4.1^43^. Biosynthetic gene clusters including RiPP peptides are predicted using antiSMASH v7.1.0 ^44^. Cleavage peptides are predicted on input proteins from each genome that are <50AAs with DeepPeptide v1.0 ^45^. Small ORFS (smORFs) are predicted using a modified version of SmORFinder v1.0.0 ^46^, with modifications described below. Finally the workflow performs functional annotation with Kofamscan v1.0.0 ^47^. The user can download the entire set of KEGG HMMs or a subset for functional annotation, and can optionally run functional annotation or not. All molecule type counts and functional annotations are summarized into a main results folder. All raw files including antiSMASH genbank files, predicted smORFs and cleavage peptide sequences, and summaries of counts of each molecule type are available on Zenodo at https://zenodo.org/records/15858468.

The original SmORFinder pipeline uses a custom binary of Prodigal with a minimum gene size set to 15, and then passes predicted proteins smaller than 50 AAs for smorf prediction. In our experience some genomes will fail at the Prodigal step due to known bugs with this program. Additionally, we needed the input ORF annotation files to be uniform so that locus tags could be cross-referenced regardless of the annotation program used. Therefore we implemented an additional workflow to a fork of SmORFinder called smorfinder pre_called where pre-annotated ORFs from programs such as Pyrodigal can be directly passed to SmORFinder. The smorfinder pre_called workflow bypasses the default Prodigal step of SmORFinder and directly passes in .ffn, .ffa, and .gff files. All downstream steps are the same for smORF prediction. We give the user the option to use the default SmORFinder workflow that uses Prodigal for gene calling, or our custom pre_called workflow using ORF predictions directly from Pyrodigal. This custom fork of SmORFinder is available at https://github.com/elizabethmcd/SmORFinder where we automatically run tests on new code incorporations on a set of five test genomes. This distribution of SmORFinder is available as a Docker image that we incorporate into the workflow. In our experience because we do not implement the same minimum gene size in Pyrodigal there will be fewer smORFs predicted with the pre_called workflow, but we found this trade-off feasible in our case since we wanted to directly input ORFs predicted with Pyrodigal in order to match up annotations from different programs.

We decided to pre-filter input proteomes for proteins <50 AAs before predicting cleavage peptides with DeepPeptide for several reasons. First, in a test of running the ∼1300 species-representative genomes through the bac-mining workflow with and without the size pre-filter before predicting cleavage peptides, we did not observe a significantly higher amount of predicted cleavage peptides without the pre-filter step (33,478 cleavage peptides predicted without the filter versus 30,293 predicted with the filter). We did observe a significantly higher amount of propeptides predicted with and without the filter (220,405 propeptides predicted without the filter versus 35,349 predicted with the filter). However, propeptides are considered to be biologically inactive and we filter out these peptides for any downstream analyses such as summary statistics and bioactivity predictions. Additionally, the cost to predict cleavage peptides with DeepPeptide on entire proteomes versus those filtered to less than 50 AAs is more than 10X the amount, which we did not see it worthwhile to spend for the entire ∼11,500 bacterial genomes dataset when the increase in predicted cleavage peptides was not significant.

Additionally a small number of peptides are classified as both a cleavage peptide and smORF. In our analysis of the 1350 species-representative genomes, approximately 10% of peptides that are classified as smORFs are also classified as a cleavage peptide. Interestingly, 12% of peptides classified as an smORF are also classified as a propeptide by DeepPeptide. For counting peptide types in Figure 3B, we discard any propeptides, unless they are classified as a smORF. However we don’t toss out or distinguish between smORFs that are also classified as a cleavage peptide. For reporting bioactivity summaries below, we do apply a sequence similarity filter of 95% to reduce redundancy.

### Peptide bioactivity prediction workflow

We then input the collection of predicted peptides from the bac-mining workflow (all smORFs, cleavage peptides, and RiPP core sequences) into a Nextflow workflow for peptide-bioactivity-prediction available on GitHub at https://github.com/MicrocosmFoods/peptide-bioactivity-prediction. The peptide-bioactivity-prediction workflow takes in a set of peptide sequences and compares them to a database of known peptide sequences of the user’s choice, predicts signal peptides, physicochemical signatures, and bioactivities using machine learning classification models. The database of peptides can be any set of sequences the user inputs, such as those with known bioactivities or from a specific source or database. We created a custom database that combines peptide sequences from the FermFooDB database with sequences specifically from fermented foods and the Peptipedia database that contains sequences with experimentally tested and predicted bioactivities.

Peptide sequences can be input as a single FASTA or several FASTA files to the workflow, and the workflow will first handle combining all input sequences. Input sequences are compared to the custom database using DIAMOND blast-p v2.1.7 ^48^. Signal peptide sequences are predicted with DeepSig v1.2.5^49^. Peptide physicochemical properties such as hydrophobicity, isoelectric point etc. are predicted with the peptides.py package v0.3. Peptide bioactivities are predicted using AutoPeptideML v0.2.9^50^, a command-line and web-based tool for creating peptide bioactivity classification models and using those models for downstream predictions. We applied the 16 models made as part of the AutoPeptideML publication as well as constructing three of our own models using this tool, as described below. The models we ran as part of the AutoPeptideML release are available on Zenodo https://zenodo.org/records/13363975 and include bioactivity predictions for antibacterial, ACE inhibitor, anticancer, antifungal, antimalarial, antimicrobial, antioxidant, antiparasitic, antiviral, blood-brain barrier crossing, DPPIV inhibitor (also known as gliptins, they act on incretin hormones such as GLP-1 and gastric inhibitory peptide which maintain glucose homeostasis by increasing insulin secretion and decreasing glucagon secretion), anti-MRSA, neuropeptide, quorum sensing, toxic, and tumor T-cell antigens.

### Peptide bioactivity machine learning models

We were interested in predicting the anti-inflammatory potential of the predicted peptides, since recent clinical research has shown that fermented foods may interact with human receptors involved in these activities^11^. We chose to construct models using the AutoPeptideML tool since this is an open source and user-friendly tool for reproducibly creating peptide bioactivity models. For each of these models, we pulled sequences from the Peptipedia database labelled as anti-inflammatory^51^. We filtered to only sequences where predicted = False, since we only wanted to include sequences for building and evaluating the models that had some amount of experimental or literature data to support that label. This resulted in a total of 3903 sequences for the anti-inflammatory model. Using the AutoPeptideML v.1 web server^52^, we uploaded the TSVs of curated, labelled peptides for both bioactivities. We selected “automatic search for negative peptides” and for building the anti-inflammatory model we selected immunological as an overlapping bioactivity. We kept the homology partitioning threshold at the default of 0.3 and using the default alignment algorithm as mmseqs. Scripts documenting how the sequences were filtered are available on GitHub: https://github.com/MicrocosmFoods/peptide-model-curation and all model files including configs, raw sequences, and results files are available on Zenodo: https://zenodo.org/records/16749254. Performance metrics for each model as reported in the output of AutoPeptideML are listed in Table 1 below.

### Peptidomics data curation and bioactivity prediction

We collected 5 proteomics datasets that had data available either through the PRIDE database or in supplementary tables. This included proteomics experiments from sheep milk kefir ^27^, four different cheeses ^24^, fermented cocoa beans ^28^, donkey milk ^29^, and kefir grains ^30^. Scripts for generating FASTA files from either the PRIDE Xml table deposits or supplementary tables are available on GitHub at https://github.com/MicrocosmFoods/fermented-food-peptidomics-mining-results. Each set of peptides from each sample was predicted for bioactivity and physicochemical properties using the peptide-bioactivity-predictor workflow. Notably, the raw sequences were not pre-processed or deduplicated in any way, so there are many sequences that are highly similar to each other.

## Supplementary Figures and Tables

**Supplementary Figure 1.**
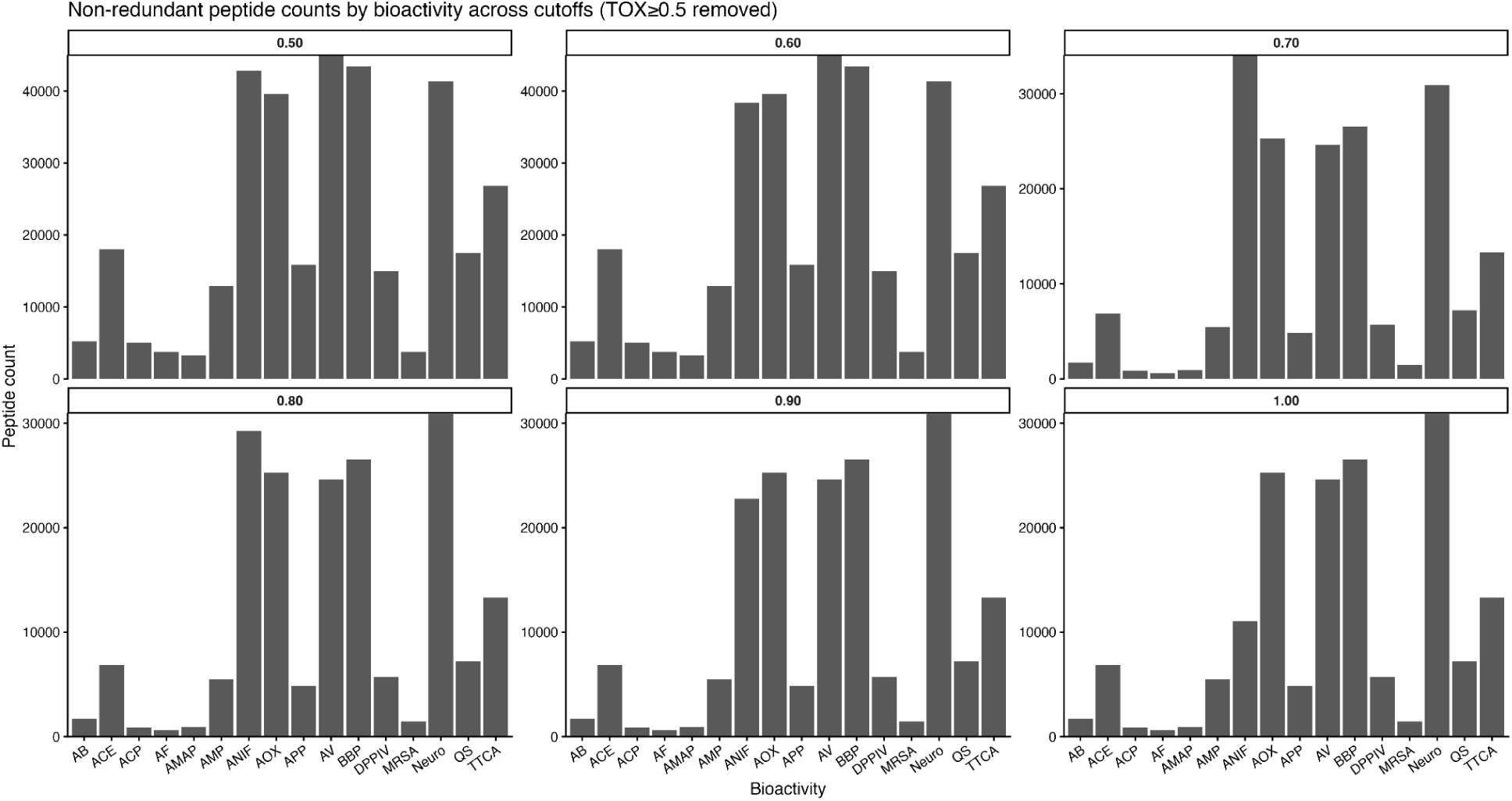
Comparisons of probability threshold cutoffs for peptide bioactivity predictions for genome-encoded peptides. Each facet plot demonstrates the probability threshold cutoff to show how many non-redundant, non-toxic peptides (defined as probability above 0.5) remain within that probability threshold.

**Supplementary Figure 2.**
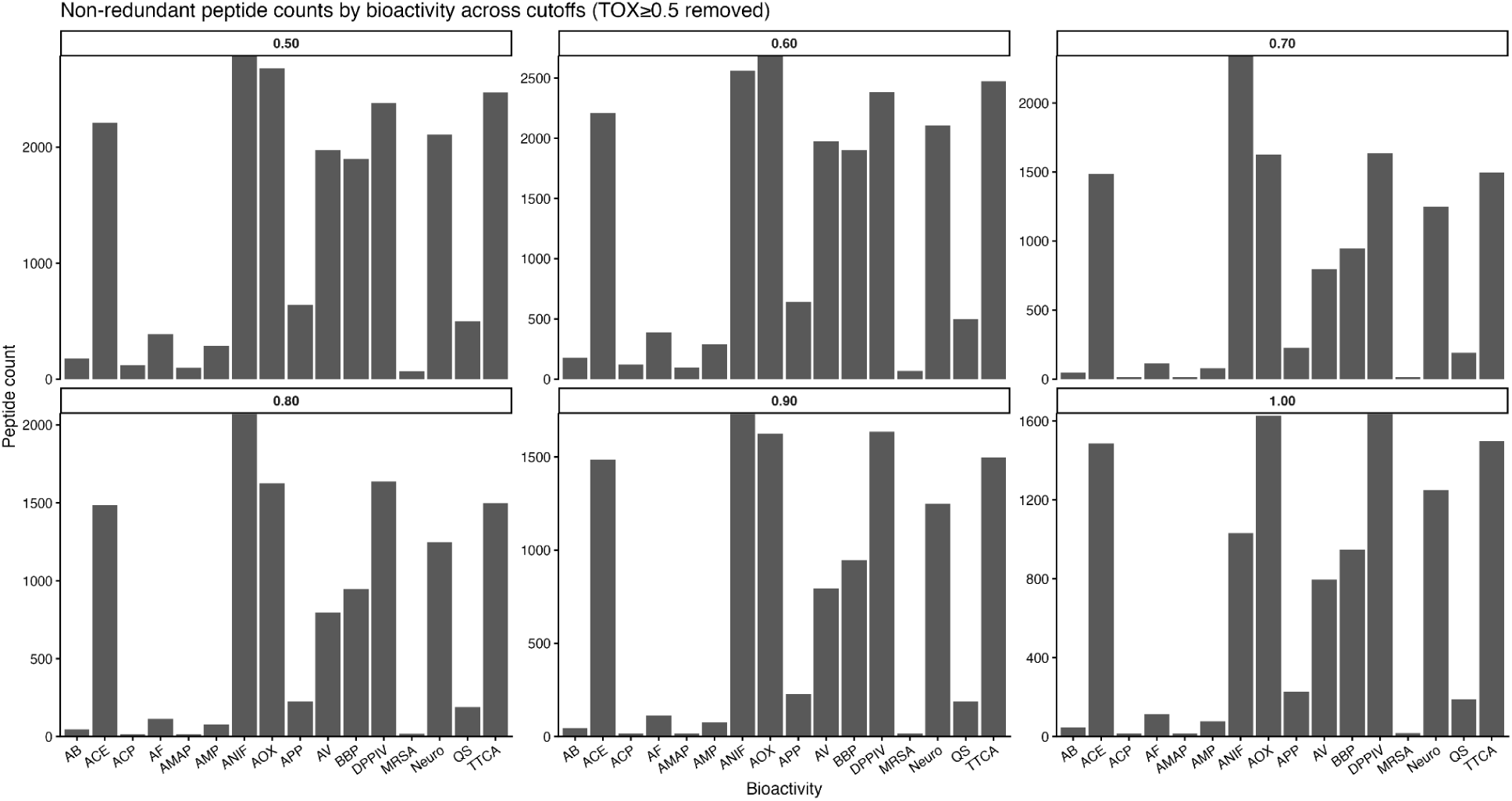
Comparisons of probability threshold cutoffs for peptide bioactivity predictions for proteomics-sourced peptides. Each facet plot demonstrates the probability threshold cutoff to show how many non-redundant, non-toxic peptides (defined as probability above 0.5) remain within that probability threshold.

**Supplementary Table 1.**
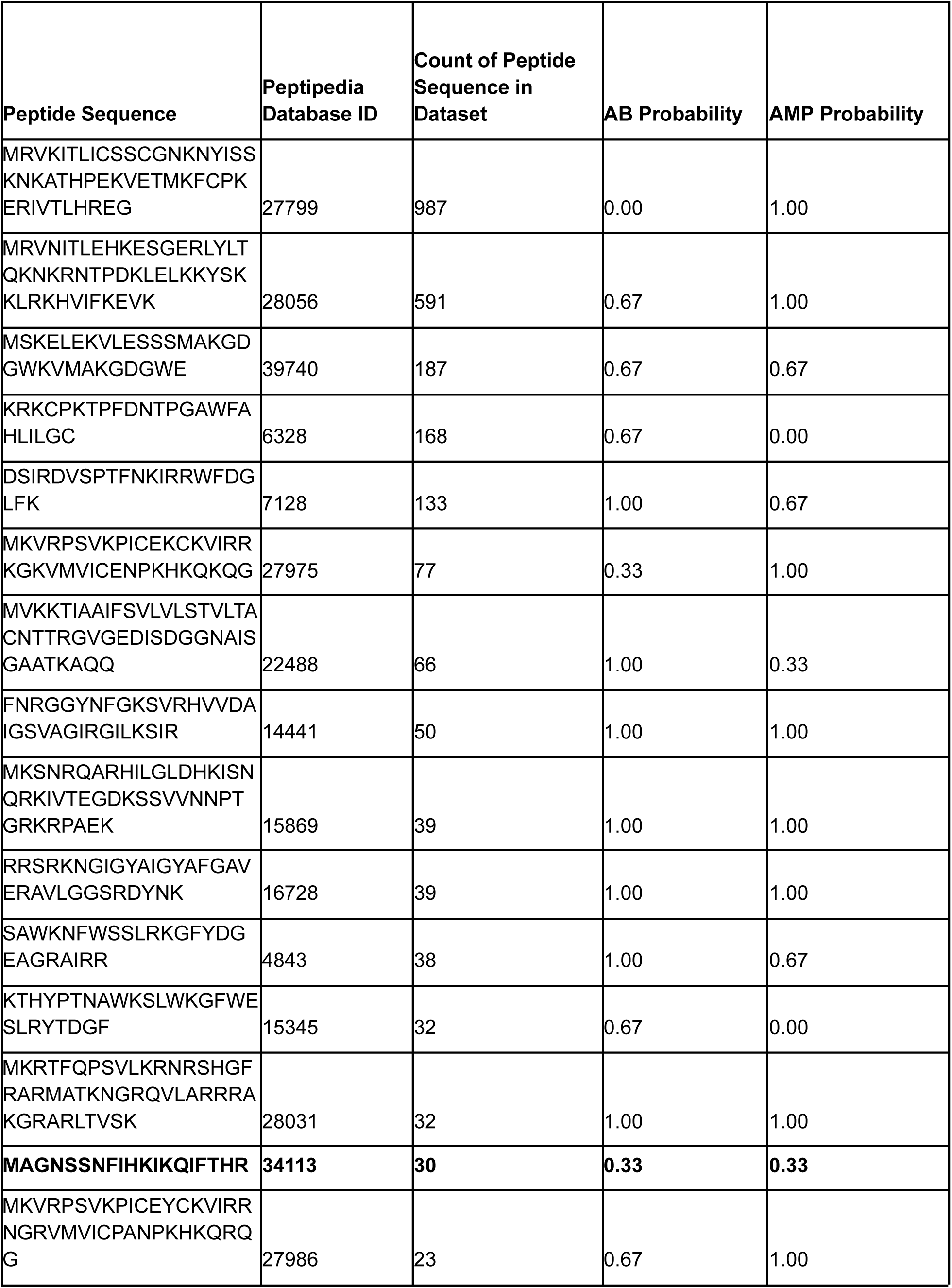
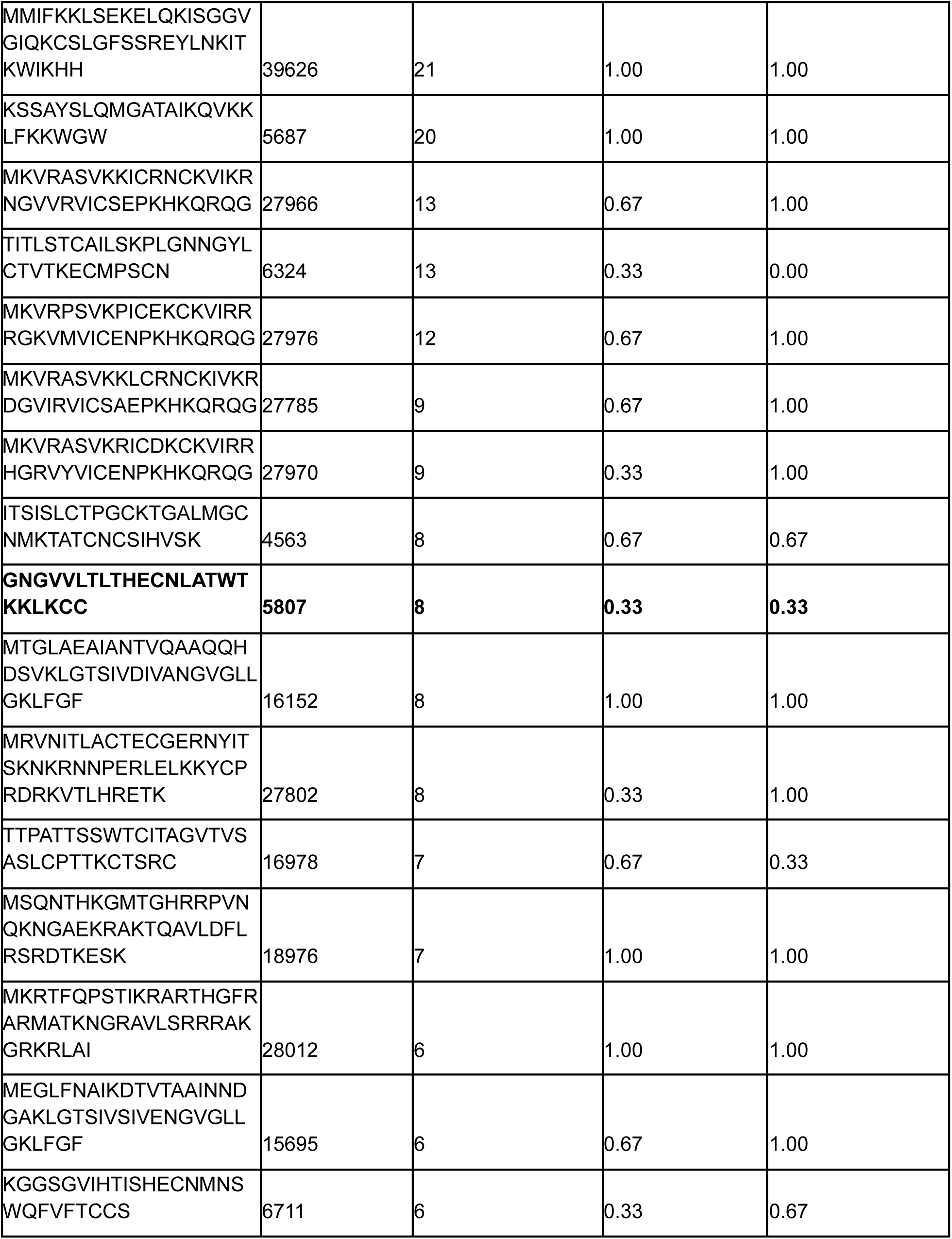
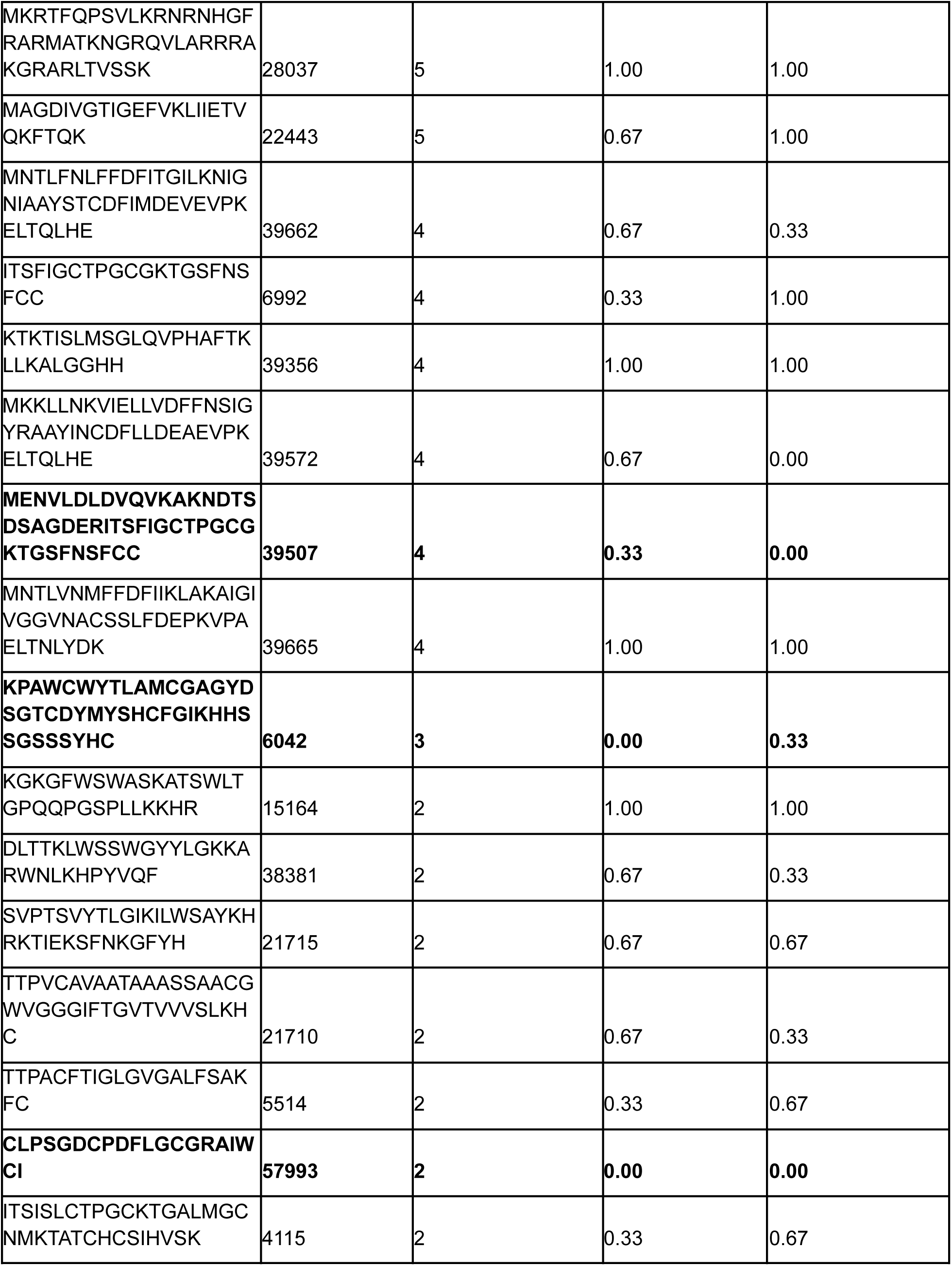
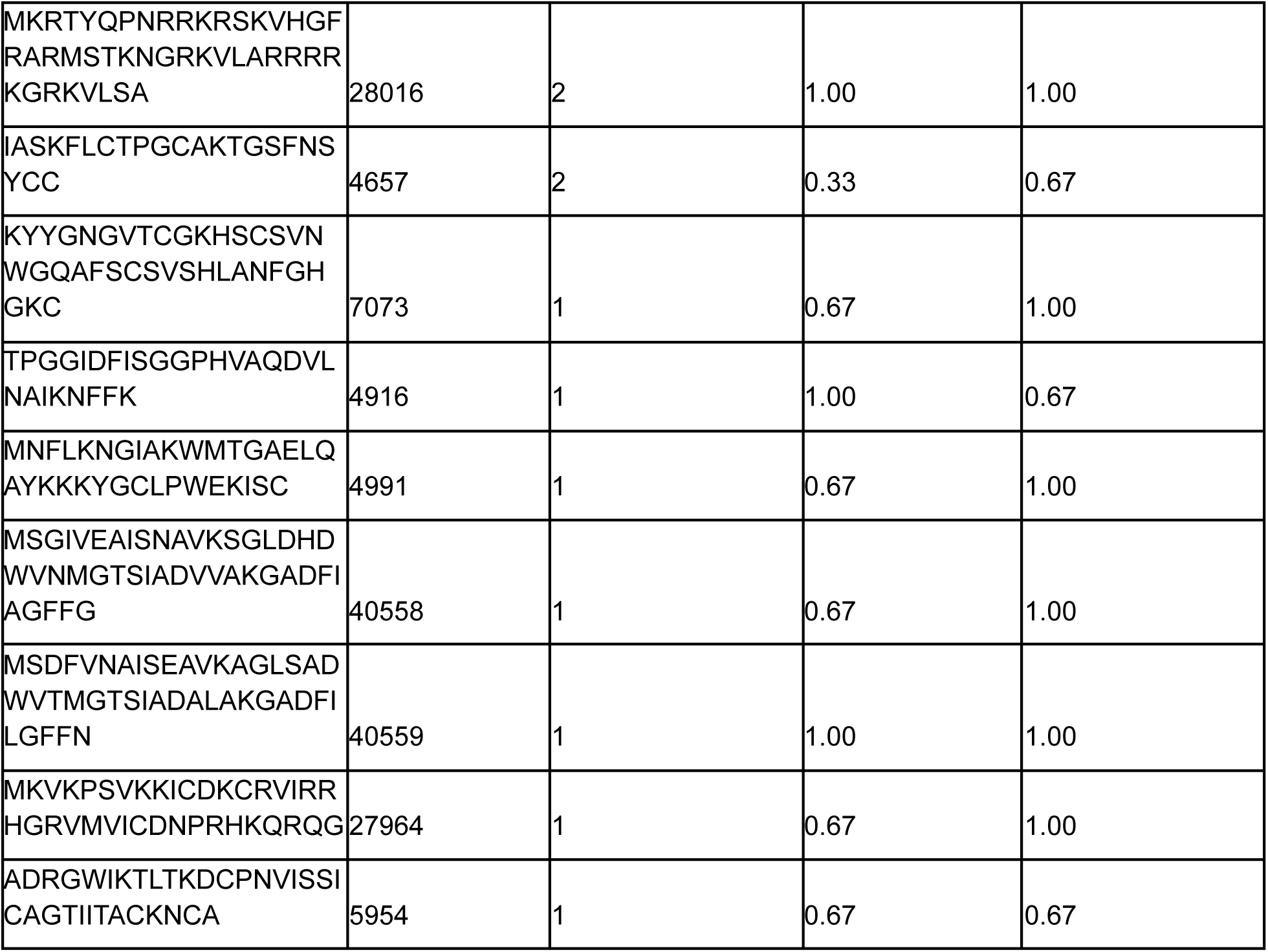
Genome-Encoded Peptides with Hits to Sequences in Peptipedia with Experimentally Verified Antibacterial Activity and Machine Learning Model Probabilities. For genome-encoded peptides, we performed DIAMOND blast-p searches to peptides with experimentally validated antibiotic activity in the Peptipedia database. We then pulled sequence hits that had 100% sequence similarity over the exact length match of the peptide. The peptide count is the count of that exact peptide sequence in the genome-encoded dataset. Machine learning model probabilities for the antibiotic (AB) and antimicrobial peptide (AMP) models are reported for the sequence in the dataset. Peptides where neither model has a predicted probability < 0.5 are highlighted in bold.

**Supplementary Table 2.** Peptides from Fermented Food Proteomics Experiments with Hits to Sequences in Peptipedia with Experimentally Verified Anti-inflammatory Activity and Machine Learning Model Probability. For peptides from multiple proteomics experiments, we performed DIAMOND blast-p searches to peptides with experimentally validated anti-inflammatory activity in the Peptipedia database. We then pulled sequence hits that had 100% sequence similarity over the exact length match of the peptide. The peptide count is the count of that exact peptide sequence in the proteomics datasets. Machine learning model probabilities for the anti-inflammatory (ANIF) model is reported for the sequence in the dataset.

| Peptide Sequence | Peptipedia Database ID | Peptide Count | ANIF Model Probability |
| --- | --- | --- | --- |
| QLLRLKKYKVPQLEIV<br>PN | 10222 | 38 | 1.0 |
| **YLGYLEQLLRLKKYK**<br>**VPQ** | **10426** | **2** | **0.47** |
| IHAQQKEPMIGVNQE<br>LAY | 9976 | 2 | 1.0 |
| EPMIGVNQELAYFYP<br>ELF | 9825 | 2 | 1.0 |

**Supplementary Table 3.**
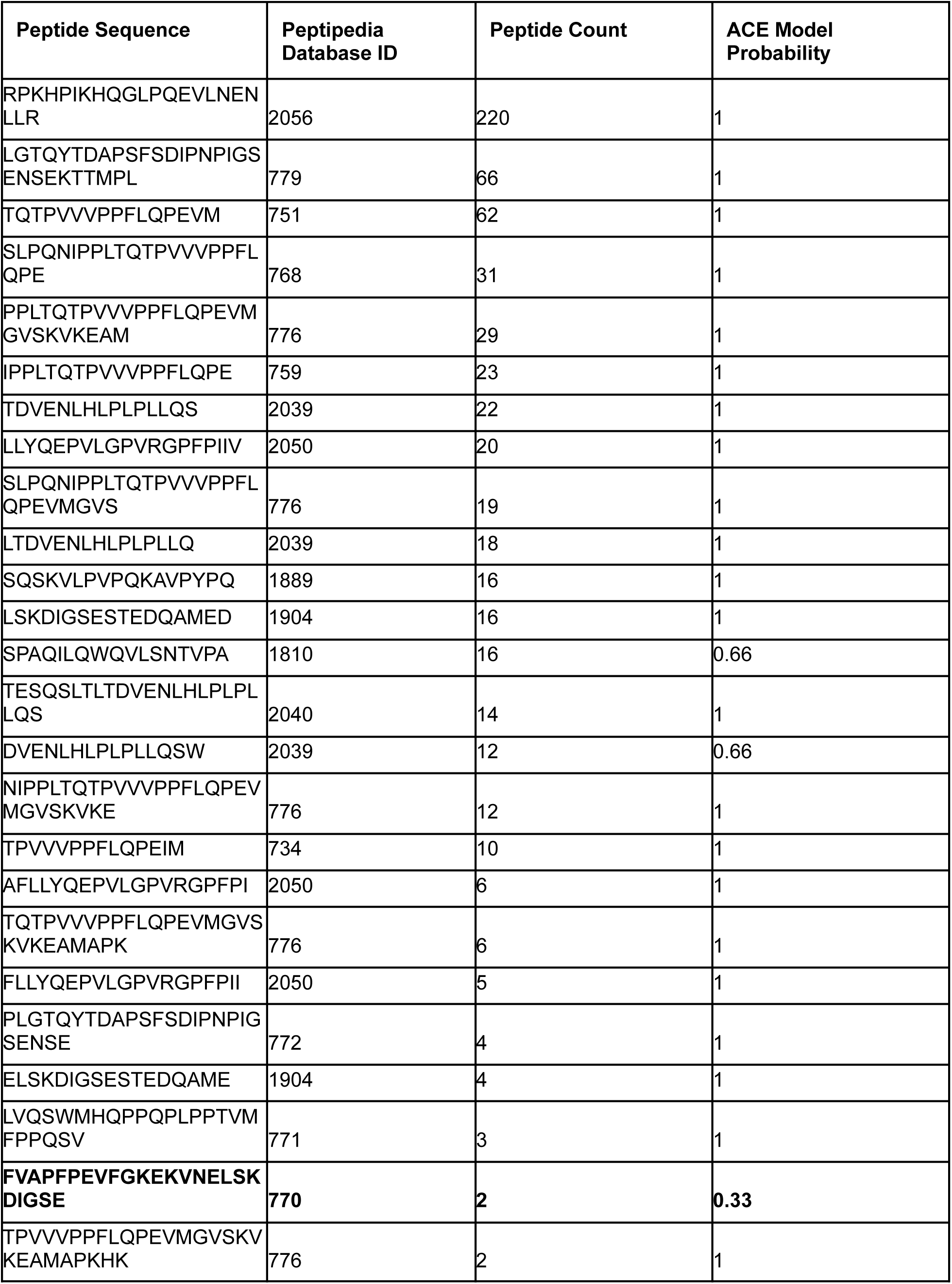
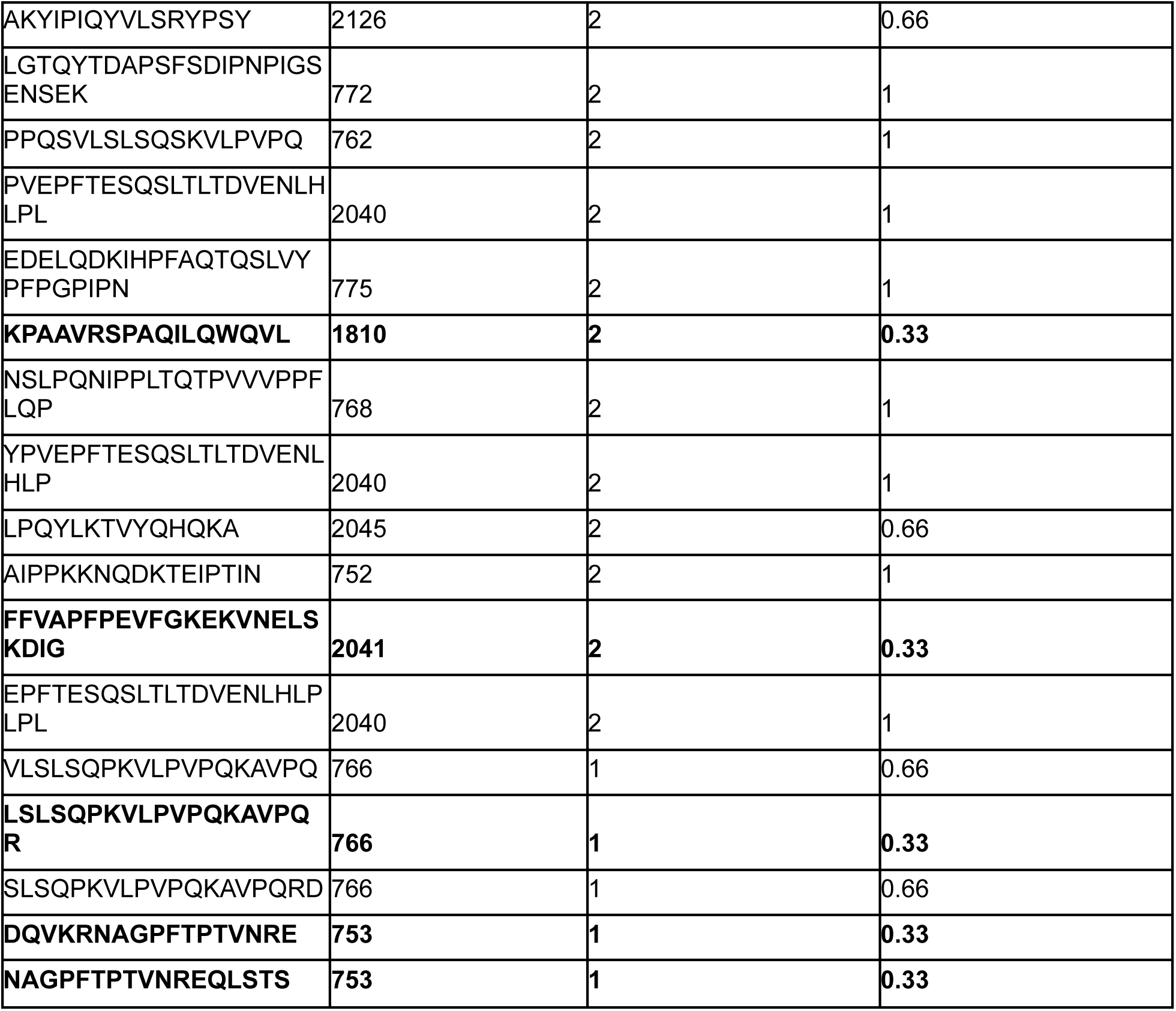
Peptides from Fermented Food Proteomics Experiments with Hits to Sequences in Peptipedia with Experimentally Verified ACE-Inhibitor Activity and Machine Learning Model Probability. For peptides from multiple proteomics experiments, we performed DIAMOND blast-p searches to peptides with experimentally validated anti-inflammatory activity in the Peptipedia database. We then pulled sequence hits that had 100% sequence similarity over the exact length match of the peptide. The peptide count is the count of that exact peptide sequence in the proteomics datasets. Machine learning model probabilities for the ACE-inhibitor (ACE) model is reported for the sequence in the dataset. Peptides where the model has a predicted probability < 0.5 are highlighted in bold.

## Acknowledgements

We would like to thank Elisa Caffrey for consulting on construction of the microbial genomes database and advice on what would be useful to the community. We thank Ray Keren for assisting in metadata curation of the microbial genomes database. This research was funded by the Astera Institute as part of Rachel Dutton’s residency project.

